# Evolution from adherent to suspension – systems biology of HEK293 cell line development

**DOI:** 10.1101/2020.01.29.924894

**Authors:** Magdalena Malm, Rasool Saghaleyni, Magnus Lundqvist, Marco Giudici, Veronique Chotteau, Raymond Field, Paul Varley, Diane Hatton, Luigi Grassi, Thomas Svensson, Mathias Uhlen, Jens Nielsen, Johan Rockberg

## Abstract

The need for new safe and efficacious therapies has led to an increased focus on biologics produced in mammalian cells. The human cell line HEK293 has bio-synthetic potential for human-like production and is today used for manufacturing of several therapeutic proteins and viral vectors. Despite this increasing popularity there is still limited knowledge of the detailed genetic and composition of derivatives of this strain. Here we present a genomic, transcriptomic and metabolic gene analysis of six of the most widely used HEK293 cell lines. Changes in gene copy and expression between industrial progeny cell lines and the original HEK293 were associated with cellular component organization, cell motility and cell adhesion. Changes in gene expression between adherent and suspension derivatives highlighted switching in cholesterol biosynthesis and expression of five key genes (RARG, ID1, ZIC1, LOX and DHRS3), a pattern validated in 63 human adherent or suspension cell lines of other origin.

## Introduction

The production of protein therapeutics is a fast-growing field as it allows for the generation of sophisticated molecules with high specificity and activity in humans (Leader et al., 2008)(Bandaranayake and Almo, 2014)(Alex Philippidis, 2017; Walsh, 2014)(Alex Philippidis, 2017; Walsh, 2014). The need for proper protein folding and glycosylation of therapeutic proteins has promoted their production in mammalian cells, especially Chinese hamster ovary (CHO), but also human cells such as HEK293 (Zhu et al., 2012)(Dumont et al., 2016)(Priola et al., 2016; Sanchez-Garcia et al., 2016; Walsh, 2014). Even though CHO is the most common mammalian cell factory used for production of advanced recombinant proteins, there is an increasing demand for improved and more efficient bioproduction platforms. With an increasing number of difficult-to-express proteins including next-generation biologics, such as bispecific antibodies and antibodydrug conjugates, entering clinical development, alternative or engineered expression hosts are being explored. For instance, extensive omics profiling of the CHO cell has been carried out during recent years (Wlaschin et al., 2005)(Hammond et al., 2011)(Xu et al., 2011)(Brinkrolf et al., 2013)(Birzele et al., 2010)(Becker et al., 2011)(Sellick et al., 2011)(Dietmair et al., 2012), which has paved the way for cell line engineering efforts aiming to improve bioproduction efficiency and product quality (Xiao et al., 2014)(Lee et al., 2015)(Kildegaard et al., 2013). Moreover, human production cell lines, such as HEK293, can serve as convenient expression hosts for proteins with specific requirement for human post-translational modifications (Dumont et al., 2016)(Lalonde and Durocher, 2017).

The human cell line HEK293 is the most commonly utilized human cell line for expression of recombinant proteins for various research applications. This cell line originated from the kidney of an aborted human female embryo and was originally immortalized in 1973 by the integration of a 4 kbp adenoviral 5 (Ad5) genome fragment including the genes encoding the E1A and E1B proteins, at chromosome 19 (Graham et al., 1977) (Louis et al., 1997). The expression of E1A and E1B enable continuous culturing of HEK293 cells by inhibiting apoptosis and interfering with transcription and cell cycle control pathways (Berk, 2005). In addition, since E1A and E1B are essential helper factors for adeno associated virus (AAV) production, the continuous expression of these genes makes HEK293 cells attractive production hosts for recombinant AAV particles (Clément and Grieger, 2016).. The genomes and transcriptomic profiles of six different HEK293 cell lines were recently mapped, confirming previous observations of a pseudotriploid genome with the adenoviral DNA inserted on chromosome 19 (Y. C. Lin et al., 2014)(Bylund et al., 2004)(Louis et al., 1997). Furthermore, it was shown that standard laboratory culturing of the cell lines in most cases did not change the overall genomic composition of the cells although the organization of the HEK293 genome is continuously evolving through the events of chromosomal translocations and copy number alterations (Y. C. Lin et al., 2014). However, the chromosome number of HEK293 cells derived from different commercial sources has been reported to show a high degree of variation, suggesting that long-term cultivation and subcloning of cells result in karyotypic drift (A.A. Stepanenko and Dmitrenko, 2015). Such abnormalities and genomic instability is, however, characteristic for immortalized cells and have also been reported for CHO cells (A.A. Stepanenko and Dmitrenko, 2015)(Väremo et al., 2014)(Vcelar et al., 2018)(Wurm, 2013).

Several HEK293 cell lines have been established from the parental HEK293 lineage. Strategies to improve recombinant protein expression include the generation of transformed HEK293 cells constitutively expressing the temperature sensitive allele of the large T antigen of Simian virus 40 (293T), (DuBridge et al., 1987) or the Epstein-Barr virus nuclear antigen EBNA1 (293E) (Murphy et al., 1992)(Swirski et al., 1992). Cell lines expressing these viral elements, enable episomal replication of plasmids containing the SV40 origin of replication (293T) or EBV oriP (293E). One 293E cell line is host for the production of dulaglutide (TRULICITY^®^)(Lalonde and Durocher, 2017). In addition, several HEK293 cell lines have been adapted to suspension growth in serum-free medium at high cell density (Graham, 1987)(Garnier et al., 1994)(Côté et al., 1998), enabling largescale cultivation and bioproduction in bioreactors. Two industrially relevant suspension cell lines are 293-F and 293-H (Gibco, Thermo Fisher Scientific). The 293-F clone was initially isolated by limiting dilution of HEK293 where a clone with fast growth and high transfectivity was obtained, which was later recloned and subjected to suspension growth adaptation in serum-free medium (**Figure 1A**). Interestingly, the 293-H cell line is a suspension growth adapted clone originally derived from a more adherent HEK293 cell clone. The final 293-H clone has high adherence during plaque assays, fast growth, high transfectivity and productivity (**Figure 1A**). The 293-F cell line is today used for the production of the regulatory approved rhFVIII (NUWIQ^®^), whereas 293-H cells are used for production of rFVIIIFc (ELOCTATE^®^) and rFIXFc (ALPROLIX^®^)(Dumont et al., 2016)(Lalonde and Durocher, 2017). Furthermore, the Freestyle 293-F cell line has been generated by adaptation of 293-F cells to Freestyle medium (Gibco, Thermo Fisher Scientific). Even though each of the above-mentioned cell lines are all derived from the same original HEK293 strain, significant genomic and transcriptomic changes between parental and progenitor cell lines can be expected due to the genomic instability of HEK293 as discussed above.

**Figure 1.**
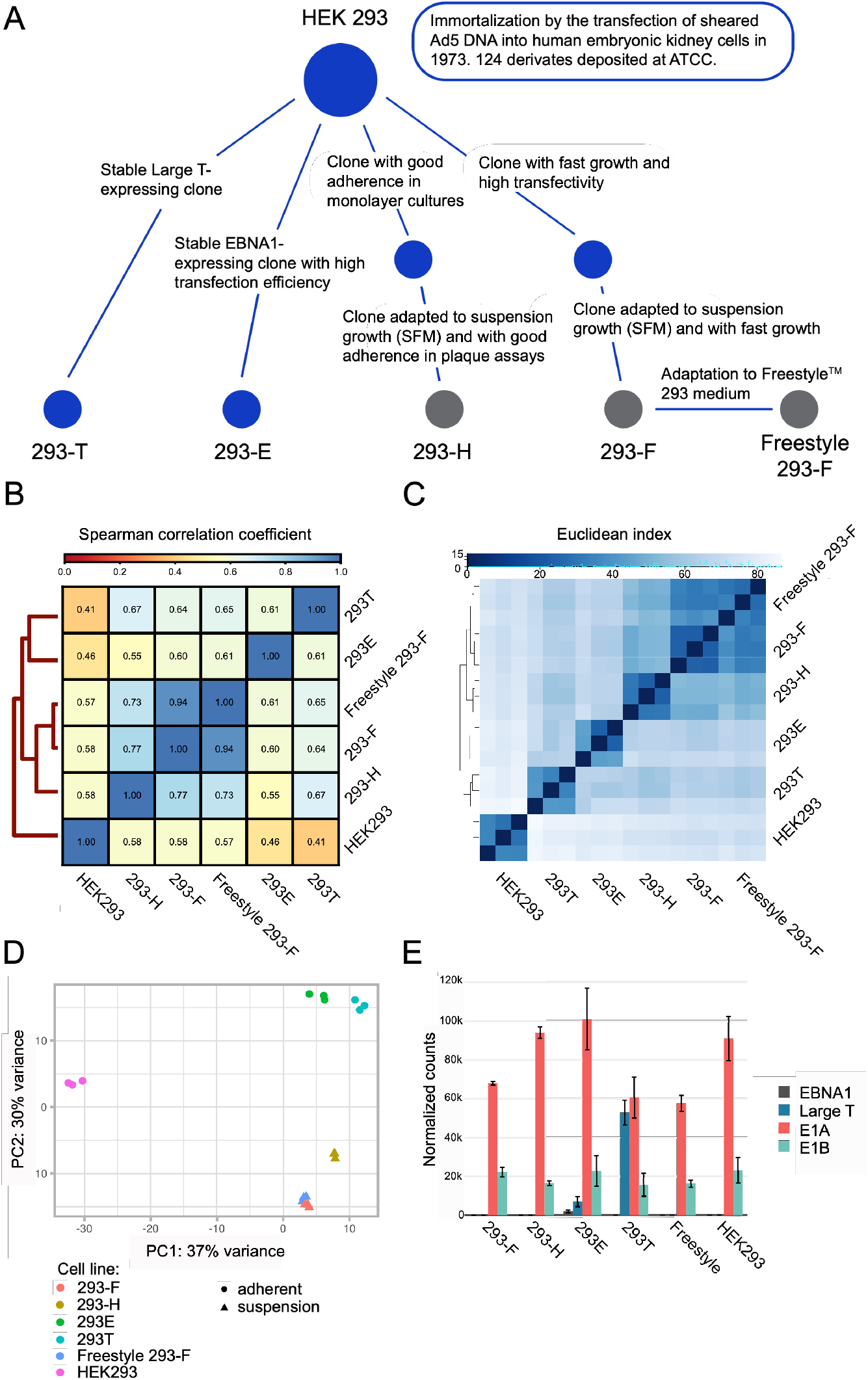
Comparisons of genomic and transcriptomic profiles of HEK293 cells showed taxonomic divergence between parental HEK293 and progeny cell lines. (A) A schematic overview of the lineage relationship of the six HEK293 cell lines used in this study. Blue dots represent adherent cells whereas grey dots represent suspension cell lines. (B) Genomic comparison between HEK293 cell lines based on Spearman correlation coefficients of read counts. Darker blue color indicates higher correlation. (C) Sample-to-sample comparison between transcriptomes illustrated by a heatmap and hierarchical clustering of taxonomical divergence between samples. Darker blue color indicates shorter Euclidean distance between samples and more similarity. (D) PCA plot showing the separation in expression pattern between samples. (E) RNA expression levels of stably integrated viral genes (EBNA-1, Large T, E1A and E1B) in various HEK293 cell lineages determined by RNA sequencing.

Despite extensive usage of CHO and HEK in both suspension and adherent mode and several empirical protocols for adaptation in either direction, molecular knowledge of the key genes involved in the transition between the two growth states is limited. While adherent cells have traditionally been widely used for the production of viruses, e.g. AAV and lenti virus for clinical research, suspension cells are the platform of choice for the bioproduction of therapeutic proteins. The main advantages of suspension cell bioproduction schemes are growth at very high cell densities in serum-free medium at scale, enabling higher volumetric yields. Easy adaptation between cellular growth in adherent to suspension mode has traditionally been a key feature when selecting strains for bioprocess applications, including usage of CHO and HEK293. Whereas certain experimental steps are more efficient in adherent mode, e.g. chemical transfection and viral infection, the ability to increase the volumetric cell density by growth in suspension without cell clump formation that results in oxygen limitations is a key step from a manufacturing perspective.

Here, we present a genomic and transcriptomic analysis of the HEK293 parental cell line along with five widely used HEK293 derivatives (**Figure 1A**). An overall analysis of the differences in genomic landscape and transcriptomic profiles was performed in order to provide novel molecular insights into the differences between cell lines that have occurred during the process of clonal isolation and expansion. Furthermore, we focus on transcriptomic differences between adherent and suspension HEK293 cells and the impact of the differentially expressed genes on metabolic pathways and the phenotype of the cells from a bioprocess perspective.

## Results

### Overall genomic and transcriptomic profiles emphasize clonal divergence between all the progeny cell lines compared to the parental HEK293 cell line

In this study, six industrially relevant HEK293 cell lines (**Figure 1A**) were subjected to omics profiling. This set of cell lines includes the parental HEK293 as well as five additional cell lines that have all been clonally derived from parental HEK293 cells. The cell lines can be divided into either adherent (HEK293, 293E and 293T) or suspension (293-H, 293-F and Freestyle 293-F) cells. The genomes and the transcriptomes of these six cell lines were sequenced using Illumina HiSeq. **Table S1** provides full results of transcript levels (TPM) for all cell lines. Comparisons of the genomes and transcription profiles between the cell lines show overall similar results (**Figure 1B and 1C**). Hierarchical clustering divided the progeny cell lines into two different taxonomic groups, of either adherent (293T, 293E) or suspension cell lines (293-H, 293-F and Freestyle 293-F), diverged from the parental HEK293. Interestingly, the original HEK293 cell line was the most distant from all other cell lines. As expected, the two 293-F lineages (293-F and Freestyle 293-F) showed very similar profiles. The same pattern of gene expression clustering was visualized by principal component analysis (**Figure 1D**), where the suspension cell-lines grouped together in the plot, with a very close clustering of 293-F and Freestyle 293-F cells. On the other hand, the adherent cell-lines HEK293E and HEK293T showed larger variations in gene expression patterns between cell lines. The parental cell line HEK293 showed a notable difference in transcriptome profile compared to all the other cell lines along the first principal component (PC1). These results indicate a genomic divergence of the clonal lineages compared to the parental HEK293 and suggest the presence of similar transcriptomic traits between HEK293 progeny cell lines individually selected for during the isolation of each clone. Hierarchial clustering of the cell lines based on SNPs gave a similar trend with high similarity between 293-F and Freestyle 293-F cell lines, whereas the parental HEK293 cell line had lower similarities to the progeny the cell lines (**Figure S1A**). However, a different pattern of overall clustering was observed, with the original HEK293 and 293E cell lines separated from the rest on a separate branch and the lowest similarity scores observed for the 293E cell line compared to the rest.

The HEK293 cell line was originally immortalized by the random integration of viral genomic DNA of adenovirus 5 (Graham et al., 1977), which includes the E1A and E1B genes. In this study, overall high mRNA levels of E1A and E1B were observed in all HEK293 cell lines (**Figure 1E**), with significantly (p<0.05) higher expression levels of E1A in HEK293, 293E and 293-H cell lines over the other cell lines, whereas 293F had a significantly (p<0.05) higher E1B expression compared to 293H and Freestyle 293-F (**Figure S1B**). No other significant differences were observed for E1B. Furthermore, the 293T and 293E cell lines were originally generated by the transfection and stable integration of plasmids expressing the SV40 Large T antigen and EBNA-1, respectively. As expected, the gene expression of Large T and EBNA-1 was detected in 293T and 293E, respectively (**Figure 1E**). Interestingly, expression of the Large T antigen was also observed in 293E, which is not reported by the supplier (ATCC). The presence of a truncated version of Large T in the 293E genome was confirmed by de novo assembly of all reads not mapping to the human reference genome (**Figure S1**). Tracing the origin of the 293E cell line (Swirski et al., 1992), the Large T expression of 293E may be derived from the pRSVneo plasmid that was used to cotransfect HEK293 cells along with the pCMV-EBNA plasmid for the generation of the stable EBNA-1 expressing clone (293c18) by G418 selection. The pRSVneo plasmid contains a truncated version of the Large T gene (according to the AddGene vector Database), which aligns perfectly with the truncated Large T sequence found in the 293E genome (**Figure S1C**).

### Copy number variation between the parental HEK293 cell line and its derivatives confirmed the genomic instability of the HEK293 strain and identified loci with conserved copy number gain/loss between progeny cell lines

In order to evaluate the genomic variation between HEK293 and its derivatives further, overall genomic copy number variation of all progeny cell lines compared to the parental HEK293 was performed. A comparison of gained and lost regions on all chromosomes between all cell lines can be found in **Figure 2** and **Table S2**. Interestingly, a conserved pattern of copy number gain or loss of large regions has occurred on several chromosomes of all HEK293 progeny cells compared to HEK293, whereas other changes are more local or cell line specific. For instance, on chromosome 13, a region of >15 Mb (**Figure 2**) has been amplified significantly in all cell lines compared to the parental HEK293 strain. All elements with copy number gain of >1 log2 fold-change common to all progenitor cells are located in this region (**Table S2**). Amongst these, four out of seven protein-coding genes (BORA, MZT1, PIBF1 and KLHL1) belong to the cytoskeleton gene set (GO:0005856). On chromosome 18, there is a conserved pattern of copy number loss of most of the chromosome sequence for all progeny cell lines compared to the parental HEK293 strain, with the exception of a high degree of copy number gain of a region close to the centromere for all cell lines except 293E. Within the region of conserved gain, several genes encoding cell adhesion molecules within the desmocollin (DSC) and desmoglein (DSG) subfamilies, belonging to the cell-cell adhesion gene set (GO: 0098609), are located. When analyzing more local copy number variations between progeny cell lines and the parental strain, some interesting loss or gain of full or partial elements compared to the parental HEK293 was identified. For instance, copy number loss was observed for the fumarate hydratase (FH) gene, which has previously been reported to have lost several gene copies in HEK293 and hence been hypothesized to play a role in the phenotypic transformation of HEK293 (Y.-C. Lin et al., 2014). Interestingly, the fumarate hydratase gene along with the neighboring kynurenine 3-monooxygenase (KMO) gene, had a log2-fold copy ratio of <−1 in 293T, 293-F and Freestyle 293-F cell lines compared to the parental HEK293 (**Figure 2** and **Figure S2A**), suggesting that these cells have half the number of copies compared to the parental cell line. Moreover, the 293T and 293-H cell lines have a gain of the genomic loci surrounding the FH gene, while maintaining the copy number of the FH gene compared to HEK293. Interestingly, the resulting FH expression levels of the cell lines only partly correlated with the gene copy number changes (**Figure 2** and **Figure S2A**). Even though the gene copy number of the parental HEK293 strain is the same as for 293T and 293-H lineages, the FH mRNA levels of HEK293 was as low as the expression levels of the lineages with only half number of FH gene copies. Moreover, the expression levels of KMO was comparably low in all cell lines but did not correlate with gene copy number. Besides the changes in gene copy number of the FH loci, a loci around the transducin-like enhancer protein 4 (TLE4) gene, encoding a transcriptional co-repressor of Wnt signaling pathway members, and the non-coding RNA LINC01507 was found to have a log2-fold copy number gain of >1.5 in all progeny cell lines except for 293E (**Figure 2)**. This gain in the TLE4 loci was accordingly reflected in the transcription level of the gene with a higher level of expression in 293T, 293-H, 293-F and Freestyle 293-F compared to 293E and HEK293 (**Figure S2A**). In addition, a major loss of copy number of the ADAM3A pseudogene was observed for all cell lines accept 293-H with a maintained low or no expression of the psuedogene observed in the cell lines (**Figure 2** and **Figure S2AB**).

**Figure 2.**
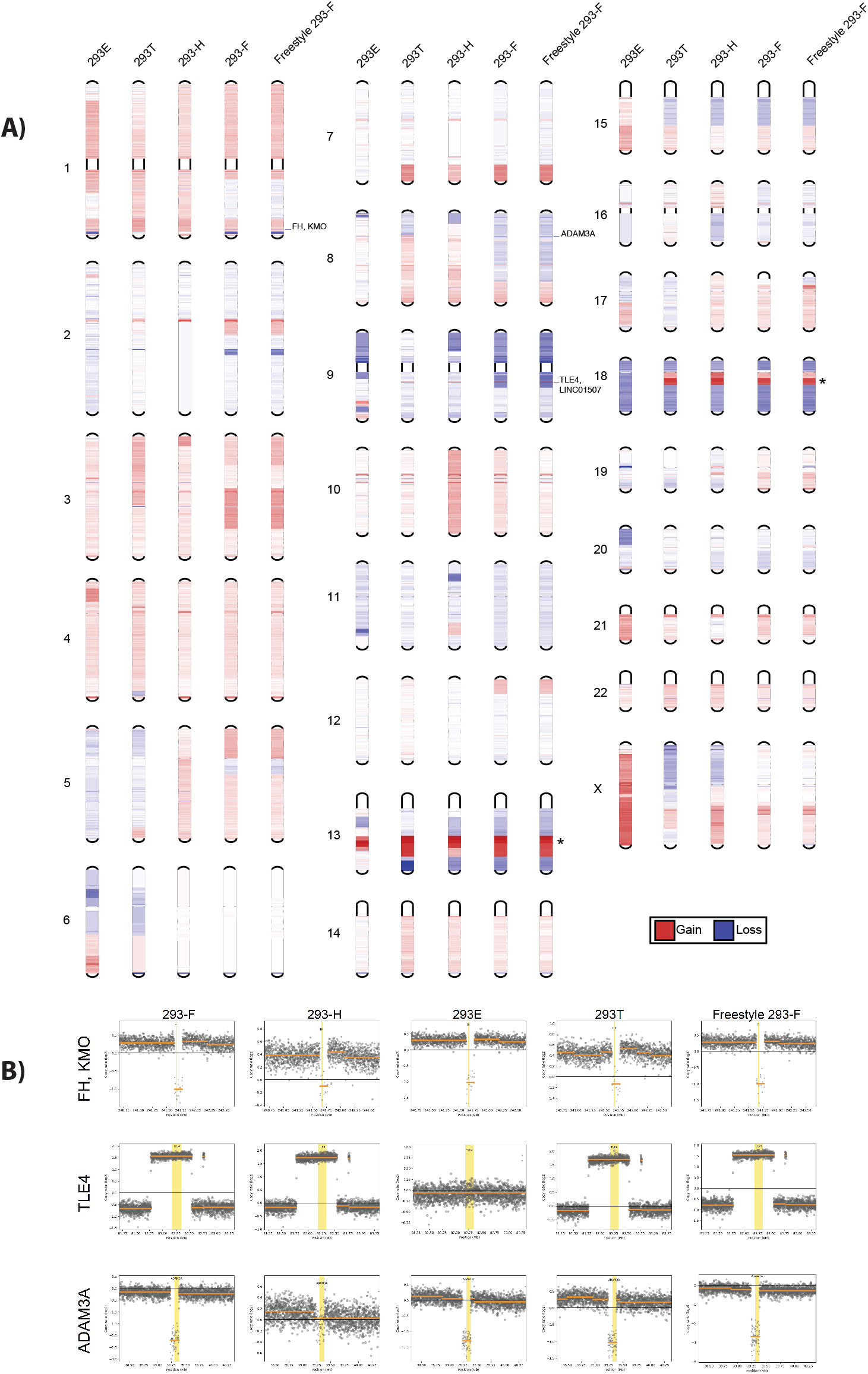
Copy number variation analysis of HEK293 progeny cells compared to the parental HEK293 revealed conserved patterns of copy number gain and loss. **A)** Copy number gain (red) or loss (blue) over all chromosomes of progeny HEK293 cell lines. The color intensiveness correspond to log2 foldchange compared to HEK293 cells. The asterix symbol (*) marks regions of common copy number gain (>1 log2 fold change) common between several progeny HEK293 cell lines on chromsome 13 and 18. Identified genes with local copy number gain or loss of the full gene sequence common between several HEK293 progeny cell lines are marked with the gene name. B) Copy number gain or loss (log2 fold change compared to HEK293 for each cell line of the FH, KMO,TLE4 and ADAM3A genes.

Due to the observed pattern of common genomic changes to progeny cell lines compared to the parental HEK293, an evaluation of common SNPs amongst all progeny cell lines but not HEK293 was performed. GO enrichment analysis of common genes with high or moderate impact SNPs different in all progeny cell lines compared to the original HEK293 (**Table S3**), showed significant (adjusted p-value <0.05) enrichment of homophilic cell adhesion via plasma membrane adhesion molecules (GO:0007156; adjusted p-value 0.025; fold enrichment 10.26; data not shown) and cell-cell adhesion via plasma-membrane adhesion molecules (GO:0098742; adjusted p-value 0.032; fold enrichment 7.53; data not shown). All genes with moderate or high impact SNPs in progeny cell lines compared to HEK293 found amongst both these GO-terms were protocadherins (PCDH12, PCDHB10, PCDHB13, PCDHB15, PCDHB16 PCDHGA2, PCDHGA3 and PCDHGB2). In addition, the Teneurin-2 gene (TENM2) (within GO:0098742) also had an altered SNP allele in all progeny cell lines compared to HEK293. These SNPs all result in missense mutations with unknown biological impact on the gene products. However, the enrichment of common SNPs within this group of genes in all HEK293 progeny cell lines may suggest an impact on the protein function and a selective advantage of such phenotypic changes during continuous cell line cultivation.

### Differential expression between all progeny cell lines and the parental HEK293 revealed consensus changes in gene expression of cellular component organization and integral components of the plasma membrane amongst progeny cell lines

Based on the overall genomic and transcriptomic profiles of the different HEK293 cell lines, the parental HEK293 strain stood out as different compared to all other cell lines. In order to evaluate common changes between all progeny cell lines and the parental HEK293, differential expression analysis was performed. Results showed a significant consensus of down-regulation of genes involved in extracellular matrix organization, locomotion and cell adhesion in progeny cells compared to the parental HEK293 strain (**Figure 3A**). Moreover, amino acid metabolism and metabolic process of small molecules were found up-regulated in all progeny cell-lines. Along with changes in extracellular matrix genes, there is also a consensus amongst progeny cell lines compared to HEK293 of differential expression of genes involved in other types of cellular component organization such as cell morphogenesis, cytoskeleton organization, membrane organization and cell junction organization. A comparison between gene expression fold changes and copy number variation of the differentially expressed genes (log2-fold change >+/− 1) for each progeny cell line compared to HEK293 showed a trend of enrichment of gained gene copies amongst genes with up-regulated mRNA levels (**Figure S2B**). However, there was not a clear trend of loss in gene copy number amongst transcriptionally down-regulated genes for all cell lines.

**Figure 3.**
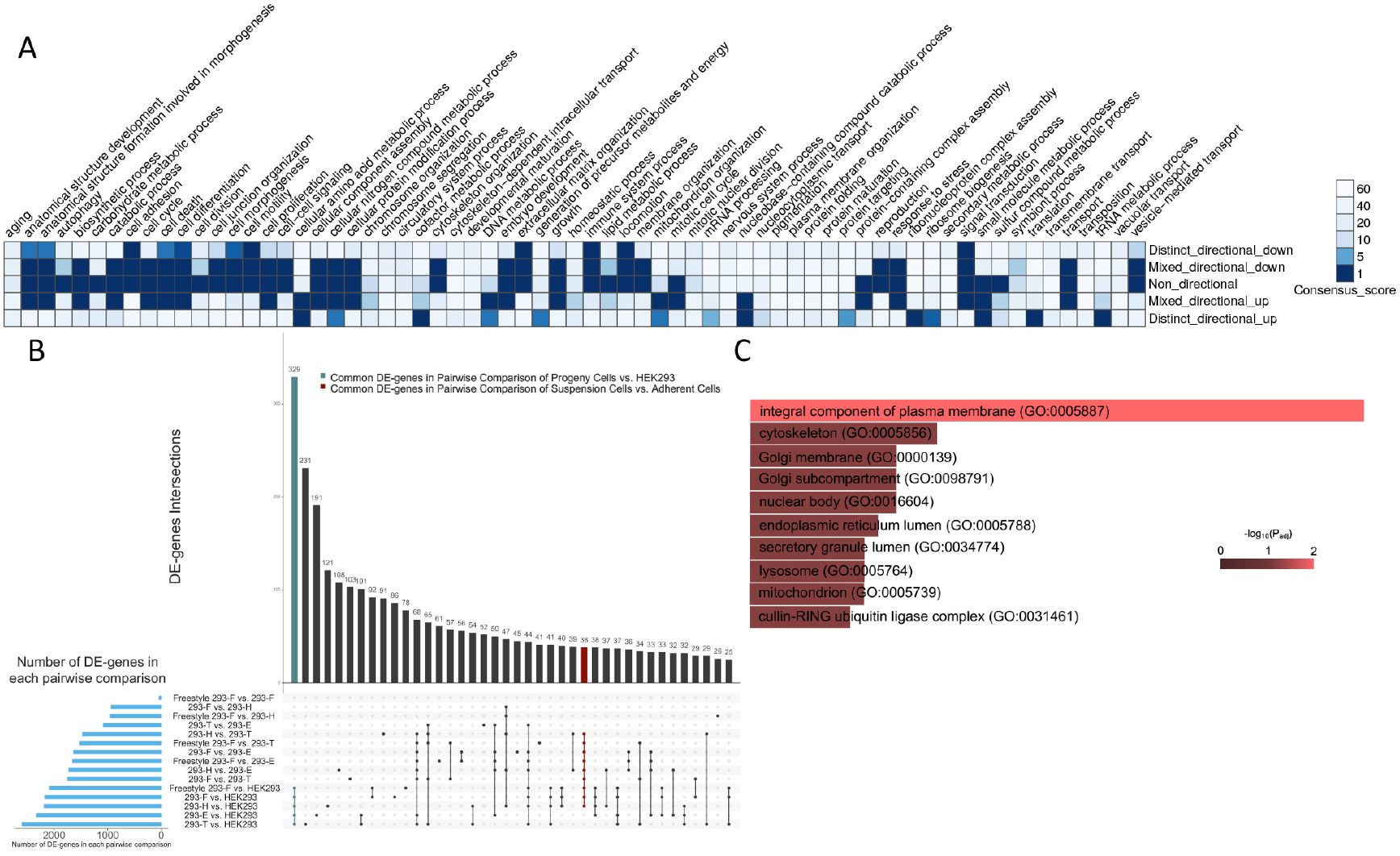
Differential expression analysis emphasized processes and genes with common changes in all progeny cell lines compared to the parental HEK293. (A) Consensus heatmap of GO biological processes with a different expression pattern between progeny cell lines compared to the parental HEK293 (B) Common differentially expressed (DE) genes in pairwise comparisons of all HEK293 cells. Blue bars show number of DE genes in each pairwise comparison. Green bar shows 329 common DE genes in pairwise comparisons of progeny with HEK293 parental cells. Red bar shows common 38 DE genes in the comparison of suspension cells against adherent cells. (C) Top ten significant GO cellular components of the 329 common DE genes in pairwise comparisons between progeny cells and HEK293.

For further evaluation of the transcriptomic similarities and changes between HEK293 cell lines, pairwise differential expression comparisons between all cell lines were performed. As expected, the parental cell line had the highest number of differentially expressed genes when compared to all other cell lines (**Figure 3B** and **Table S4**). In addition, when looking at differentially expressed genes unique to certain comparisons, the largest group of genes were found common to all pairwise comparisons between HEK293 and each of the progeny cell lines (green bar in **Figure 3B**), again emphasizing a relatively high degree of common transcriptomic changes amongst progeny cell lines differentiated from the parental HEK293. As the progeny cell lines had an enrichment of differentially expressed genes associated with cellular component organization compared to HEK293, we sought to evaluate to what cellular compartments the 329 genes common to all pairwise comparisons between HEK293 and progeny cell lines localize. In line with the overall differential expression evaluation (**Figure 3A**), which emphasized changes in for instance cell adhesion and extracellular matrix organization, there was a significant (padj <0.05) enrichment of genes relating to the integral compartment of plasma membrane (GO:0005887) amongst the common differentially expressed (DE) genes unique to the comparisons between HEK293 and all progeny cell lines (**Figure 3C**).

### Differential expression between suspension and adherent HEK293 cell lines identified key changes related to cholesterol metabolism

The growth morphology of bioproduction cell lines is of great importance for culture maintenance and efficiency of industrial bioprocessing. In order to look into gene expression variations correlating with adherent and suspension HEK 293 cell lines, differential expression analysis between adherent and suspension HEK293 progeny cell lines was performed. As results from the overall comparison of transcriptomic profiles of the HEK293 cell lines showed that the parental HEK293 cell line is highly differentiated from all of the progeny cell lines and moreover, that the Freestyle 293-F cell line is very similar to the 293-F cell line, HEK293 and Freestyle 293-F were excluded from this analysis, so as not to skew the data. Enrichment analysis of the differentially expressed genes between adherent (293T and 293E) and suspension (293-H and 293-F) progeny cell lines showed significant expression differences of similar gene sets as in the comparison between progeny cell lines and the parental HEK293 (**Figure 3A and 4A, Table S5**). For instance, the suspension progeny cell lines had a significant upregulation of gene sets involved in cellular compartment organization such as cell morphogenesis, cell junction organization, cell membrane organization and cytoskeleton organization. Interestingly, there is no significant change in the expression of the extracellular matrix organization gene set between suspension and adherent HEK293 progeny cell lines. Perhaps as expected, there was significant differential expression observed for the cell adhesion, cell differentiation, cell morphogenesis and cell motility gene sets. Interestingly, all of the abovementioned gene categories, including cell adhesion, were up-regulated in suspension HEK293 cells as compared to adherent. When looking at the most significantly differentially expressed genes (adjusted p-value <0.01) amongst the cell adhesion gene set, many genes of the cadherin superfamily of cell adhesion molecules were found up-regulated in suspension cell lines compared to adherent HEK293 progeny cells (**Table S6**).

**Figure 4.**
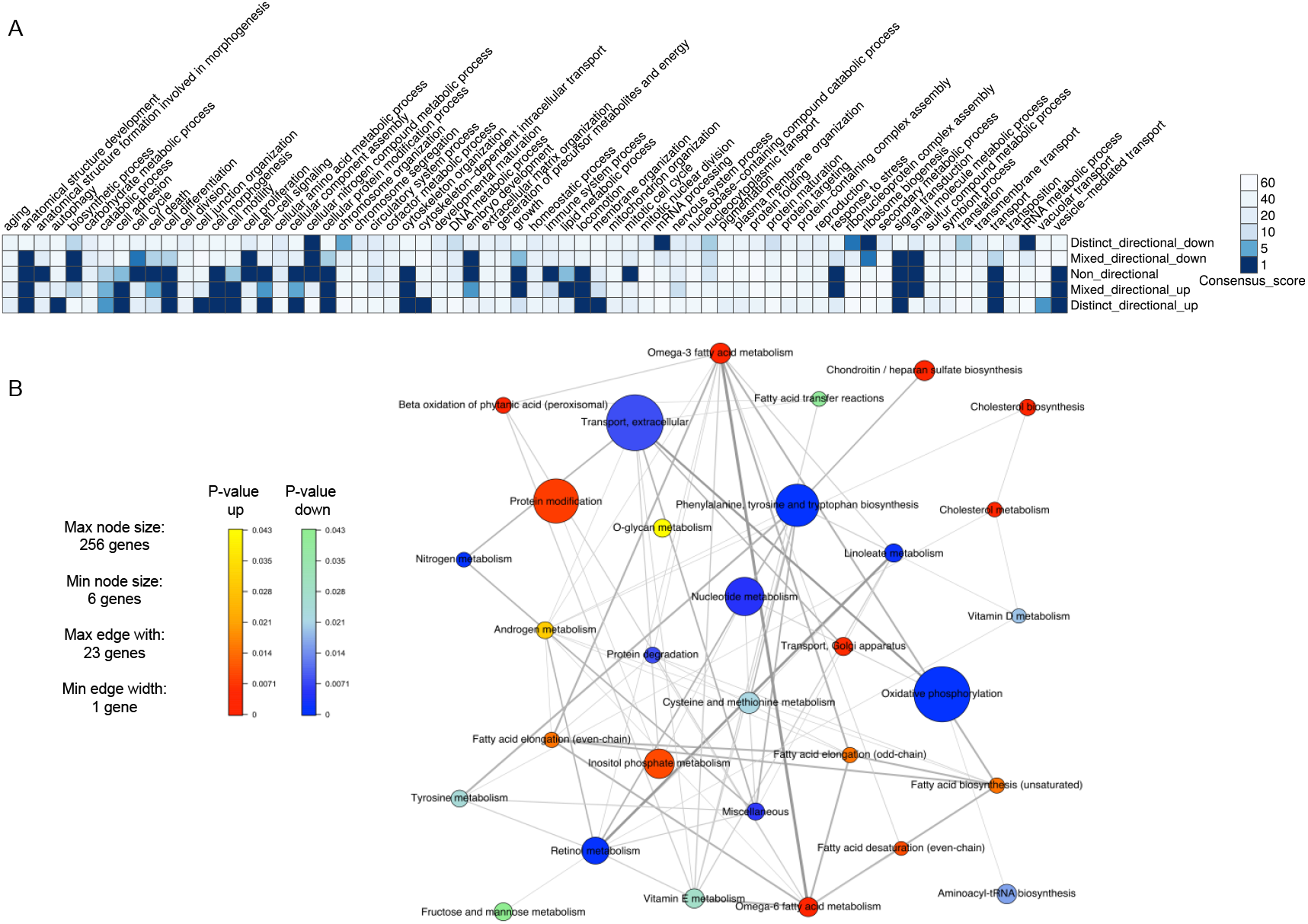
Differential expression analysis between suspension and adherent HEK293 progeny cell lineages identified significant up-regulation of gene sets associated with cell adhesion, motility, differentiation, cellular compartment organization and cholesterol metabolism in suspension cells. (A) Consensus heatmap of GO biological processes with a different expression pattern between suspension cells (293-H and 293-F) against adherent cells (293E and 293T). (B) Metabolic genes set analysis for comparing metabolic differences between suspension and adherent HEK293 progeny cell lines. The size of each node corresponds to the number of genes in each of these pathways, thickness of connections between nodes corresponds to the number shared genes between pathways and the colors of the nodes shows the p-value for the given metabolic process.

In order to evaluate what metabolic impact the differentially expressed genes may have on cells in suspension compared to the adherent state, a generic human metabolic model, HMR2 (Mardinoglu et al., 2014), was used to generate a set of metabolic genes and their assigned pathways to find metabolic pathways with altered expression between adherent and suspension HEK293 progeny cell lines. As shown in **Figure 4B**, pathways related to aromatic amino acids and oxidative phosphorylation were significantly down-regulated in suspension cells compared to adherent and had amongst the highest number of differentially expressed genes. In addition, pathways related to retinol, linoleate and nucleotide metabolism were also significantly down-regulated in suspension cell-lines. On the other hand, biosynthesis and metabolism of cholesterol were found to be most significantly up-regulated amongst metabolic pathways in suspension compared to adherent cells. In addition, pathways related to protein modification and fatty acid metabolism (omega-3/6 fatty acid metabolism, fatty acid desaturation, fatty acid biosynthesis and fatty acid elongation) were also upregulated in suspension compared to adherent HEK293 progeny cell lines. All results from the metabolic gene set analysis are provided in **Table S7**.

Focusing on the pairwise comparisons between HEK293 cell lines, 38 differentially expressed genes were identified common to all adherent to suspension pairwise comparisons (red column in **Figure 3B**, **Figure 5A**, **Table S4**). Looking into potential underlying genomic alterations of these genes, no high impact or moderate single nucleotide polymorphism (SNP) was detected in any of cell lines that could explain the differential expression (data not shown). Three of the genes (ARRDC3, HMGCS1 and PCYOX1L) had the same directional gene copy number variation (gain or loss) compared to gene expression fold-changes (up or down) of all suspension compared to adherent cells, which may at least partly explain the differential expression of these genes between the groups. Gene enrichment analysis of this set of 38 differentially expressed genes between adherent and suspension cells, performed using Enrichr (Kuleshov et al., 2016)(Chen et al., 2013), predicted the cholesterol biosynthetic process pathway as the cellular pathway most affected by this expression variation (**Figure 5B**). This result further emphasizes the differential expression between adherent and suspension cells of genes involved in the cholesterol pathway, mentioned above. Among the 38 common differentially expressed genes MSMO1, IDI1, NPC1L1, INSIG1 and HMGCS1 are directly related to cholesterol biosynthesis (GO:0006695 and/or GO:0008203, **Figure 5B**). Each of these genes had at least a two-fold increase in expression in the suspension cells compared with the adherent cell-lines. Based on these findings, we sought to predict the effect of the differentially expressed genes between adherent and suspension HEK293 cell lines on the cholesterol biosynthesis of the cells using Ingenuity pathway analysis (IPA). Although the MSMO1, IDI1 and HMGCS1 genes were all up-regulated in 293-F and 293-H compared to HEK293, the down-regulation of the lathosterol oxidase gene (SC5D), which gene product is in downstream steps of the pathway, resulted in a predicted reduction in cholesterol production in 293-F and 293-H cells compared to HEK293 (**Figure S3**). Comparisons between suspension cell lines (293-F and 293-H) and adherent progeny cell lines (293E and 293T) did not result in any predicted changes in cholesterol biosynthesis (data not shown).

**Figure 5.**
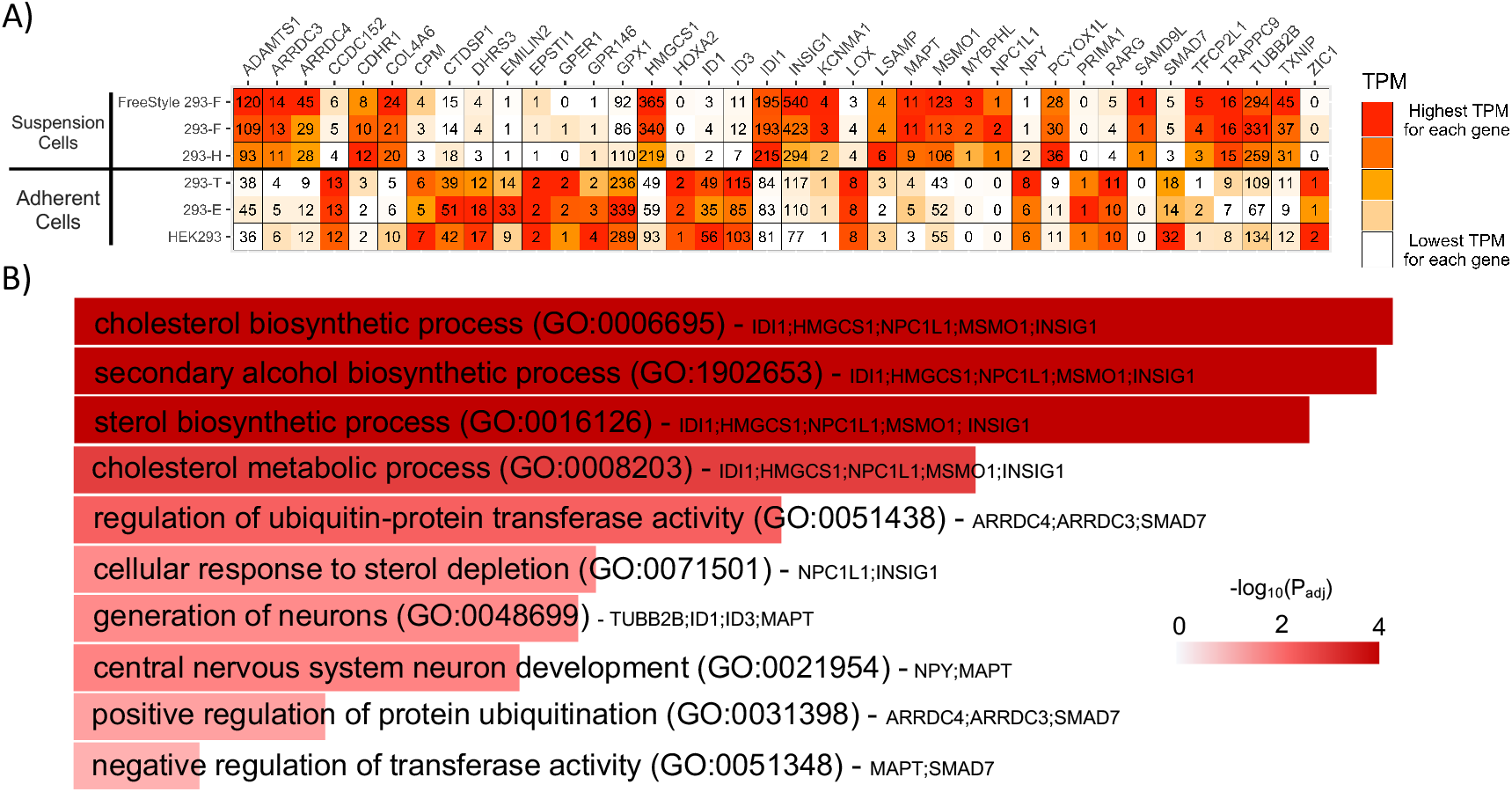
Pairwise comparison of transcription data identified cholesterol biosynthesis as the main enriched pathway between adherent and suspension HEK293 cells. (A) Heat map with TPM values for each DE gene common to all adherent to suspension comparisons. (B) The top ten most enriched biological GO terms of the 38 common DE genes between adherent and suspension cells based on gene enrichment analysis. Length and color of bars both show significance of adjusted p-value for the hypergeometric test. Also, genes mentioned in each bar are the genes that belong to enriched GO term and present in the list of 38 DE genes.

As four out of the 38 differentially expressed genes (LOX, SMAD7, ID1 & TXNIP) have previously been shown to have a role in the epithelial to mesenchymal transition (EMT) pathway (Zhao et al., 2015), we sought to evaluate the role of this pathway in the transition from adherent to suspension cell growth of these HEK293 cell lines. The normalized expression of a set of EMT markers showed that the parental HEK293 actually had the highest level of expression of various mesenchymal markers (N-cadherin, vimentin and fibronectin) of all the six cell lines (**Figure S4A**). Moreover, when predicting EMT pathway outcomes for suspension cells (293F and 293H) compared to the parental HEK293 strain using Ingenuity pathway analysis (IPA), the results predicted reduced EMT in the suspension progeny cells compared to HEK293 (**Figure S4B**). However, suspension cells were predicted to have increased disruption of adherence junctions, which is consistent with the suspension cell phenotype. Taken together the comparison between adherent and suspension HEK293 progeny cell lines suggest that the transition between adherent to suspension cell growth is not equivalent to the epithelial to mesenchymal transition even though several EMT-associated genes may be key to the difference between cell lines. Instead key changes were found associated with cholesterol biosynthesis and fatty acid metabolism.

### Extended validation of the 38 consistently differentially expressed genes between adherent and suspension HEK293 cell lines in a set of 63 human cell lines identified five key genes potentially associated with differences between adherent and suspension phenotypes

For identification of key genes involved in the transition from adherent to suspension morphology, expression data from an additional set of 63 different human cell lines deposited in the Human Protein Atlas database (Uhlen et al., 2017) were analyzed. Principal component analysis of these cell lines resulted in clustering of suspension cell lines in a distinct group separated from adherent cell lines (**Figure 6A**). However, since most of the suspension cell lines are of lymphoid or myeloid origin this clustering may be a result of a similar origin of suspension cell lines. Transcription data of the 38 previously identified differentially expressed genes from 47 adherent and 16 suspension cell lines (**Table S8**) was compared between the two groups using a Mann-Whitney U-test. Within this set, nine genes (LOX, ID1, ADAMTS1, ZIC1, KCNMA1, DHRS3, RARG, COL4A6 and ARRDC4) had significant different expression levels between adherent and suspension cell lines with p-values <0.01 (**Figure 6B and 6C**). Four of these genes (ADAMTS1, KCNMA1, COL4A6 and ARRDC4) had the opposite directional change in the extended data set compared to the differential expression between only HEK293 strains. Based on these findings, the remaining five genes (LOX, ID1, ZIC1, DHRS3 and RARG), which showed a consistent down-regulation in suspension cell lines compared to adherent cells, may play important roles in the morphological differences between the adherent and suspension cell lines.

**Figure 6.**
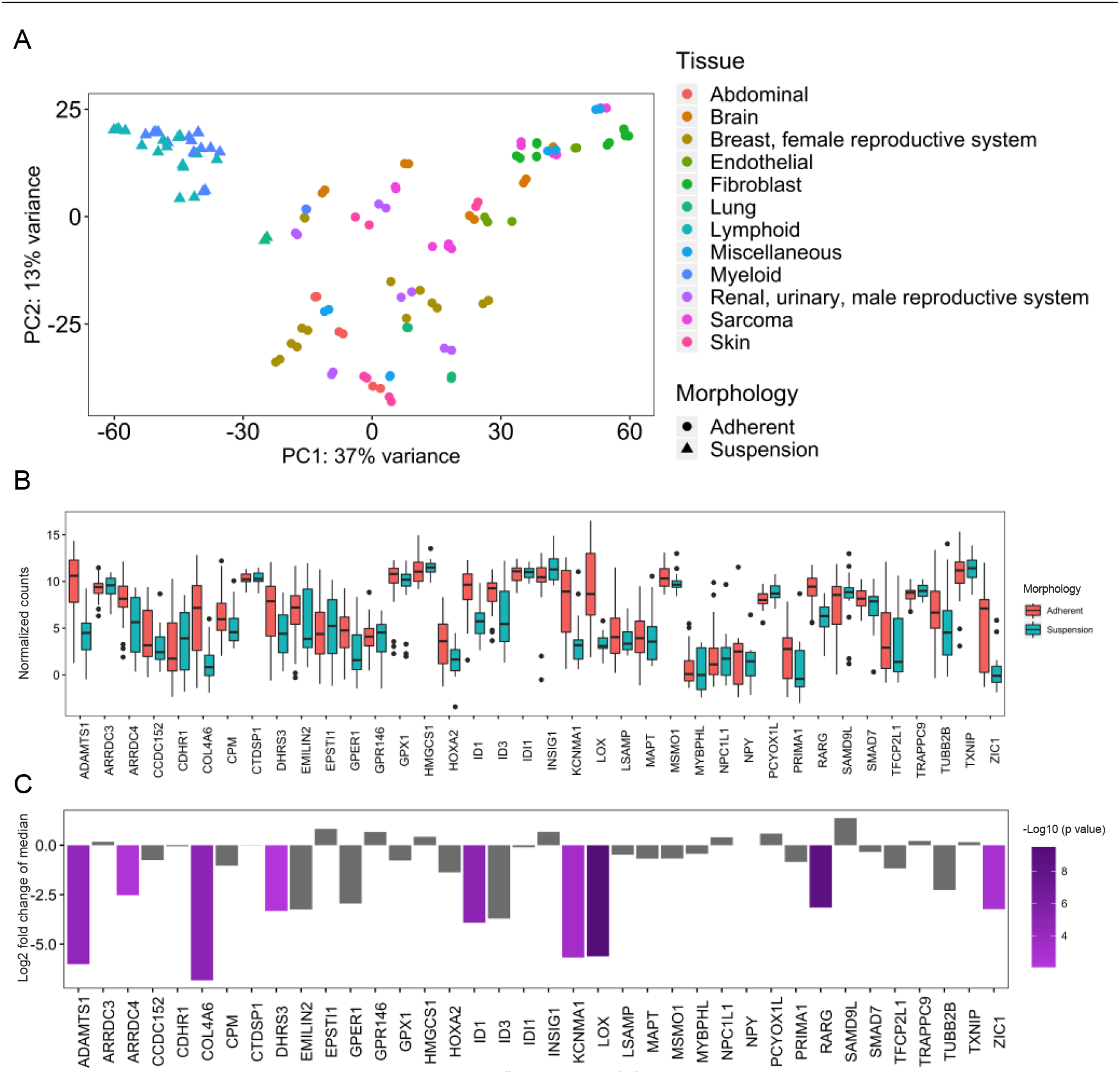
Gene expression validation of 38 previously identified differentially expressed genes in 63 human cell lines, identified nine significantly differentially expressed genes between suspension and adherent cell lines. (A) PCA transcriptomic data of 63 human cell-lines from the Human Protein Atlas shows a clear separation of suspension and adherent cell lines from different tissues. (B) Range of normalized counts in HPA cell lines for each of the previously identified 38 genes, differentially expressed between all adherent and suspension HEK293 cell lines. The black line in each box shows median of normalized counts for the gene. (C) Genes that are differentially expressed between adherent and suspension cells using a Mann-Whitney U-test, with p-values <0.01, are highlighted in purple, where length of bars shows logarithmic fold change of median between two groups and the color of bars denotes degree of significance of p-value. Non-significant genes have gray bars.

## Methods

### Cell cultivation for DNA and RNA preparation

The adherent cell lines HEK293 (ATCC-CRL-1573), HEK293T (ATCC-CRL-3216) and 293E (ATCC-CRL-10852) were obtained from ATCC and propagated in DMEM (D6429) supplemented with 10% FBS at 37°C in a humidified incubator with 5% CO_2_ in air. Suspension cell lines 293-F, 293-H and Freestyle™ 293-F (Gibco) were obtained from Thermo Fisher Scientific and cultivated in 293 SFM II medium (Gibco) supplemented with Glutamax at a final concentration of 4 mM (Gibco). Suspension cells were cultivated in 125-ml Erlenmeyer shake flasks (Corning) at 37°C and 120 rpm in a humidified incubator with 8% CO_2_ in air. All cells were propagated from frozen stocks for no longer than 20 passages.

### RNA and DNA preparation and sequencing

Adherent cells were detached by trypsinization and both adherent and suspension cells were harvested by centrifugation. Genomic DNA was extracted using the Blood and Cell Culture DNA Mini Kit (Qiagen) according to the manufacturer’s guidelines and concentrations were determined by using a NanoDrop ND-1000 spectrophotometer (Thermo Scientific). Genome sequencing was performed at the National Genomics Infrastructure (Scilifelab, Solna, Sweden) using the Illumina HiSeq X platform. For RNA extraction, cells grown in log phase from three biological replicates were collected (derived from successive propagations). Cell pellets were resuspended in RNAlater Stabilization Solution (Invitrogen) according to the manufacturer’s recommendations until RNA extraction. Total RNA was extracted from three replicates of each cell line using Qiagen’s RNeasy plus Mini Kit according to the manufacturer’s instructions. Concentrations were determined with a NanoDrop ND-1000 spectrophotometer and RNA quality was assessed on a 2100 Bioanalyzer (Agilent Technologies) using RNA 6000 Nano chips (Agilent Technologies). All samples had an RNA integrity number of at least 9.9. RNA sequencing was performed at GATC (Konstanz, Germany) using the Inview Transcriptome Advance service and an Illumina HiSeq instrument.

### DNA-sequencing analysis

Genome sequencing reads were aligned to the reference (human_g1k_v37.fasta) using bwa (0.7.12) (Li and Durbin, 2010). The raw alignments have then been deduplicated, recalibrated and cleaned using GATK (version 3.3-0-geee94ec, gatk-bundle/2.8) (McKenna et al., 2010). Quality control information was gathered using Qualimap (v2.2) (Okonechnikov et al., 2016). SNVs and indels have been called using the HaplotypeCaller. These variants were then functionally annotated using snpEff (4.1). The Piper pipeline from the National Genomics Infrastructure was used. The correlation between BAM files was assessed using multibamsummary and its plotCorrelation function from deepTools2 (Ramírez et al., 2016). Spearman was used to calculate correlation coefficients between samples,and the clusters are joined with the Nearest Point Algorithm. An SNP correlation heat map was generated using seqCAT (Fasterius and Al-Khalili Szigyarto, 2018). The heatmap was based on the similarity scores (Yu et al., 2015) between the cell lines and euclidean distances. To compare the Large T antigen sequences of 293T and 293E, unmapped reads were extracted to new bam files using SAMtools (Li et al., 2009), converted to fastq with BEDTools (Quinlan and Hall, 2010), and de novo assembled with MEGAHIT (Li et al., 2016). NCBI BLAST was used to identify the Large T antigen in the assembled contigs. To evaluate and visualize copy number variations, CNVkit (Talevich et al., 2016) was used with its whole-genome sequencing method, cbs segmentation (Olshen et al., 2011)(Venkatraman and Olshen, 2007) and the HEK293 alignment as reference. GO enrichment analysis of genes with high or moderate impact SNPs was performed using PANTHER classification system (Mi et al., 2019).

### RNA-sequencing data

Kallisto (Bray et al., 2016) was used to quantify transcripts, and the transcripts were mapped to the human GrCh37 genome. Log transformed normalized data by DESeq2 was used for cell line clustering and calculation of Euclidean sample distances. Significant testing of differential mRNA expression of E1A/B elements was done by Welsh two sample t-test. For differential expression analysis, raw count data from Kallisto was imported using the tximport package (Soneson et al., 2015) and analyzed with DESeq2 (Love et al., 2014). In the differential expression analysis, all suspension cell-lines were compared to all adherent cell-lines, and additionally, all pairwise combinations between suspension and adherent cell-lines were evaluated. The expression comparison of the viral elements was based on normalized counts from DESeq2. For evaluating differential expression of 38 common differentially expressed genes between adherent and suspension HEK293 cell lines in a set of 63 human cell lines, RNA-seq data from each cell line deposited in the HPA database was used. Based on the growth characteristics, cells were divided into two groups of adherent and suspension cells. A Mann-Whitney U-test was used to compare normalized counts based on library size between the two groups for each of the 38 differentially expressed genes.

### Gene Set analysis

To discover significant alterations of gene sets and metabolic pathways between HEK293 cell lines, the Piano package in R was used (Väremo et al., 2013). The adjusted p-values and fold changes from the differential expression was used in combination with a gene set collection based on “goslim_generic Biological Process”. The heatmap for the progeny cells lines vs. HEK293 was based on the consensus score from all pairwise gene set statistics calculations with Wilcoxon rank-sum test. The heatmap for suspension cells (293-H, 293-F) vs. adherent cells (293E, 293T) was based on the consensus score from gene set statistics calculations with mean, median, sum, stouffer and tailStrength tests. To produce the network plot, gene sets were exported from HMR2 (Mardinoglu et al., 2014). For finding differentially expressed pathways of genes between adherent and suspension cell lines, we used Wilcoxon as statistical test and filtered gene sets with adjusted p-value lower than 0.05 as significantly changed. In addition, for gene set analysis of 38 common DE genes between adherent and suspension cell lines we used EnrichR and GO biological process as gene set collection (Kuleshov et al., 2016).

### Ingenuity pathway analysis

In order to predict the pathway changes between cell lines based on differentially expressed genes from pairwise comparisons, ingenuity pathway analysis (IPA, QIAGEN Inc.,) was performed. To consider a gene as differentially expressed we used log2 fold change >1 or <1 and adjusted p-value <0.05. For filtering results of gene set analysis by IPA we used Benjamini-Hochberg multiple testing corrected p-values lower than 0.05 to find gene sets with a different expression pattern.

## Discussion

Due to the extensive usage of HEK293 cells as a bioproduction platform for pharmaceutical proteins and AAV vectors, characterization of the HEK293 genome and transcriptome is relevant for bioprocess development. A deeper knowledge of the HEK293 genomic and transcriptomic traits can for instance pave the way for more rational cell line engineering approaches, aiming to improve bioproduction efficiency and quality of protein products. As different HEK293 lineages are propagated under different conditions and the observation that immortalized continuously cultured cell lines, such as HEK293, have a high degree of genomic instability with frequent chromosome rearrangements (Y. C. Lin et al., 2014)(A. A. Stepanenko and Dmitrenko, 2015) it can be expected that different HEK293 lineages are differentiated at the genomic and transcriptomic level compared to the parental cell line. Here, the genomes and transcriptomes of the original HEK293 along with five progeny cell lineages were analyzed **(Figure 1A**). The overall comparison of genomic and transcriptomic profiles confirmed the picture of clonally diverged progeny cells as compared to the parental HEK293. A common hierarchical clustering between the cell lines was observed for both genomic and transcriptomic comparisons with the parental HEK293 cell line, the progeny suspension cell lines and the progeny adherent cells clustering on three separate branches (**Figure 1B** and **1C**). As expected, there was a high degree of genomic and transcriptomic similarities of the Freestyle 293-F and 293-F cell lines (**Figure 1B, 1C** and **1D**). It is unclear whether these two cell lines are actually the same clone or if the Freestyle 293-F cell lineage was isolated through sub-cloning of 293-F. The results presented here indeed show that they are highly similar both on a genomic and transcriptomic level and confirms the previously reported findings that standard propagation of HEK293 cell lines does not alter the genomic profile to a large extent (Y. C. Lin et al., 2014) Furthermore, based on the hierarchical clustering, the adherent progeny cell lines showed a higher degree of divergence from each other compared to suspension cells. This may be a result of the independent transformation and isolation of the 293T and 293E lineages by the stable integration of different viral genes in different labs. Whereas all the six HEK293 cell lines constitutively express the E1A and E1B genes (**Figure 1E**), which is beneficial for the production of lenti- and AAV viruses for gene therapy, the 293T and 293E cell lines also constitutively express the Large T antigen and EBNA-1, respectively (**Figure 1E**). Interestingly, a truncated version of the Large T antigen was also found to be expressed in 293E cells, which has to our knowledge not been reported previously (**Figure S1**). The Large T antigen sequence was likely derived from the pRSVneo plasmid that was co-transfected with pCMV-EBNA1 during the isolation of the 293E c18 clone (Swirski et al., 1992).

### Progeny HEK293 cell lines carry common genomic and transcriptomic traits independently evolved during cell line development

Notably, the overall comparison of the genomic and transcriptomics profiles of HEK293 cell lines suggests that the parental HEK293 strain has the highest divergence amongst the cell lines and suggests common changes independently evolved in all progeny cell lines. In line with these findings, common changes in gene copy number gain or loss (**Figure 2**) and consensus differential expression alterations (**Figure 3**) were observed amongst all progeny cell lines when compared to HEK293. Especially, a common pattern of copy number gain and loss for progeny cell lines was observed on chromosome 13 and 18 (**Figure 2**). Such patterns, found across all or several lineages of HEK293 isolated independently by different methods, suggests a selective advantage for altered copy numbers of these loci in regard to the phenotypes of the cell lines. On chromosome 13, several genes associated with the cytoskeleton (BORA, MZT1, PIBF1 and KLHL1) had a copy number gain in all progeny cell lines (**Table S2**). The DACH1 gene was also found amongst these genes, which has notably been shown to have an effect on cytoskeleton organization and to induce a more epithelial phenotype upon overexpression through the upregulation of E-cadherin(F. Zhao et al., 2015). In addition, BORA, MZT1 and DACH1 were found amongst the 329 genes commonly differentially expressed between all adherent and suspension cell lines (**Table S2**). Indeed, consensus gene expression changes associated with cytoskeleton organization was observed between all progeny HEK293 cell lines and HEK293 in the overall analysis of transcription profiles (**Figure 3A**). Within the gained region of chromosome 18 common to all progeny cell lines except 293E, there are several desmocollin and desmoglein genes (DSC1, DSC2, DSC3, DSG1, DSG2, DSG3 and DSG4) belonging to the cell-cell adhesion gene set (GO:0098609), which may render this region prone to gene copy number variation. None of these genes were found amongst the commonly differentially expressed genes between all progeny cell lines and HEK293 as may be expected since 293E had a loss of copy number in this region (**Table S4**). However, several of these genes (DSC2, DSC3 and DSG2) where found up-regulated in HEK293 suspension cells compared to adherent cells (**Table S5**), suggesting that they play a role in the differences between adherent and suspension cell lines. The observed enrichment of cell adhesion GO-terms (GO:0007156 and GO:0098742) amongst genes with common high/moderate impact SNPs unique to progeny cell lines compared to HEK293 cells also supported common genomic alterations in progeny cell lines involved in cell adhesion. Combined with the observed downregulation of the expression of cell adhesion genes in progeny cell lines compared to HEK293 (**Figure 3A**), results highlight changes in cell adhesion during continuous cultivation and cell line development of HEK293 cells.

Moreover, specific genomic regions of more local gain or loss of specific genes were observed, including a loss of fumarate hydratase (FH) gene copies. The loss of FH copies was previously observed for HEK293 by Lin and coworkers and was suggested to play an important part in the transformed phenotype of the cell line (Y. C. Lin et al., 2014). In line with this, our results showed that several of the HEK293 progeny cell lines (293E, 293-F and Freestyle 293-F) was found to only maintain half the number of FH gene copies compared to the original HEK293 (**Figure S2**), supporting an advantageous loss of the FH gene in the HEK293 cell lineages. Furthermore, a conserved pattern of substantial gain (>1.5 log2-fold change) of the TLE4 gene and surrounding loci, including the non-coding RNA LINC01507, was observed for all progeny cells except 293E (**Figure S2**). This gene encodes the transducin-like enhancer protein 4, which is a transcriptional co-repressor of members of the Wnt signaling pathway. Down-regulation of this gene has been associated with the epithelial-to-mesenchymal transition (EMT) phenotype (Wu et al., 2016), consistent with the observation in this study that the original HEK293 strain seems to have a more mesenchymal expression profile then the other cell lines (**Figure S4**). Interestingly, TLE4 has previously been reported to have both a tumor suppressor function and to be associated with promoting tumor growth in different studies of different cancers (Shin et al., 2016)(Wang et al., 2016). Moreover, a significant loss of the ADAM3A pseudogene (<−1 log2-fold change), which has previously been associated with different cancers (Barrow et al., 2011)(Liu et al., 2012), was observed in all progeny cell lines accept 293-H compared to HEK293 (**Figure S2**). The specific gain of TLE4 and/or loss of ADAM3A loci and their association with tumor development, suggest important functions of these genes in the evolution of HEK293 cell lines.

Taken together, the genomic and transcriptomic profiles of HEK293 and its progeny cell lines suggest common genomic and gene expression changes that have independently evolved in all the progeny cell lineages compared to the original HEK293, likely reflected in traits providing advantages during single cell isolation procedures. Such traits could be associated with for instance fast proliferation and maintaining evasion of normal cell senescence. Moreover, genomic and transcriptomic traits associated with for instance cell adhesion, cell motility and extracellular matrix organization may very well be under a selective pressure during single cell cloning, potentially through altered expression of cell membrane proteins. Indeed, the parental HEK293 cell line is dividing slower and is more difficult to detach by trypsinization than the other adherent cell lines. It should however be noted that the assumption that there are independently evolved common changes in progeny cell lines compared to HEK293, relies on that the HEK293 cell line used as reference in this study (ATCC-CRL 1573) is actually the very same cell line that has been used for the generation of each progeny cell line. If an older or newer version of HEK293 was used for subcloning of either progeny cell line, then the differences we see may be due to changes that occurred in this HEK293 strain only or in the process of adaptation. Due to limited literature available on progeny cell line development, we cannot completely exclude this possibility. However, we hypothesize that this cell line is, if not the same, at least highly similar to the HEK293 strain used for progeny cell line development.

### Key differences between adherent and suspension progeny HEK293 cell lines include changes in cell adhesion gene expression

In bioproduction processes for pharmaceutical proteins, suspension cell lines enable large-scale cultivation in bioreactors, which is required in order to meet the demands for marketed drugs. However, the adaptation of cells from adherent to suspension growth and the differential cultivation procedures between adherent and suspension cells induces phenotypic changes to the cell lines. In order to develop a deeper understanding of such changes, we evaluated differences in gene expression levels between adherent and suspension progeny HEK293 cells. Consensus differential expression results were found related to up-regulation of genes associated with cell component organization such as membrane, cytoskeleton and cell junction in suspension compared to adherent cells (**Figure 5A**). Moreover, significant changes were associated with cell adhesion, cell motility, cell morphogenesis and differentiation. Noteworthy, cell adhesion was found upregulated in suspension progeny cell lines and when looking into the expression of the most significant differentially expressed genes (adjusted p-value <0.01) amongst the cell adhesion gene set (**Table S6**), there are several cell adhesion molecules (CAMs) found both up- and down-regulated in suspension cells compared to adherent. Remarkably, several members of the cadherin superfamily, including many different protocadherins (PCDH), desmoglein 2 (DSG2) and desmocollin 2 (DSC2) were found significantly up-regulated in suspension cells compared to adherent cells. This family of genes are involved in the formation of adherence junctions between cells (Bruner and Derksen, 2018). Interestingly, amongst the most significant differentially expressed genes within the cell adhesion gene set, four protocadherin members showed the highest fold-change of up-regulated genes in suspension cells compared to adherent (**Table S6**). Amongst other CAMs like integrins and Ig-superfamily cell adhesion molecules (IgSF CAMs) both significantly up- and down-regulated genes were observed between adherent and suspension cell lines. The higher expression of such cell adhesion molecules in suspension cell lines compared to adherent progeny HEK293 cells may be explained by the loss of culture dish support to grow on in case of suspension cells. Upon disruption of adhesion interactions with other cells and extracellular matrix, a natural cellular response may be to increase or maintain the expression of adhesion molecules in an attempt to restore such connections. The ability of the suspension cell lines to form cell aggregates during suspension cultivation and the ease of the cells to attach to culture dish surfaces upon cultivation without shaking, can be speculated to support these findings. Such cell adhesion molecules found up-regulated in suspension cell lines may thus be appropriate cell line engineering targets for improved bioprocess performance of suspension cell lines.

### Metabolic changes of the cholesterol biosynthesis may provide support for suspension growth in serum free medium

Further evaluation of the differentially expressed genes between adherent and suspension HEK293 progeny cell lines, based on metabolic gene set analysis, highlighted metabolic differences between the cell lines. For instance, processes related to the biosynthesis of aromatic amino acids (tryptophan, phenylalanine and tyrosine) were down-regulated (**Figure 4B**) in suspension cells compared to adherent cells. Furthermore, other metabolic pathways with differences between suspension and adherent cell lines involved cholesterol biosynthesis and metabolism, omega-3 and omega-6 fatty acid biosynthesis, fatty acid biosynthesis, desaturation and elongation, retinol metabolism and linoleate metabolism (**Figure 4B**). All of these pathways are related to lipids and/or cholesterol metabolic processes. These metabolic changes may be a result of different growth media compositions used for the cultivation of either adherent or suspension cells that may imply different concentrations of for instance amino acids, glucose or serum. When reducing the number of differentially expressed genes to those that consistently showed differential expression between adherent and suspension cells in pairwise comparisons of all cell lines, the cholesterol and sterol biosynthesis and metabolism pathway were found to be most significantly different between the cell types (**Figure 5B**). As suspension cell lines are cultivated under serum free conditions, the increased expression of genes associated with for instance cholesterol in suspension cell lines may be a result of a lower cholesterol content in the medium. However, since cholesterol is a major component of the cell membrane and has an important function for membrane structure and cell signaling (Maxfield and Tabas, 2005) the differential expression of genes associated with cholesterol synthesis and metabolism may also be of importance for the different morphologies between adherent and suspension HEK293 cells. Indeed, previous studies have shown that cholesterol plays a critical role in regulating the formation of cell-to-cell interactions in endothelial cells (Corvera et al., 2000) and that depletion of cholesterol reduces cell adhesion and increases endothelial cell stiffness (Byfield et al., 2004; Norman et al., 2010). Increased cell surface stiffness has been reported for HEK293 cells in suspension compared to adherent state as a result of up-regulation and re-organization of the actin cytoskeleton (Haghparast et al., 2015). This may partly be a result of altered cholesterol levels in the cell membrane since cholesterol is a regulator of the actin cytoskeleton and cholesterol depletion has been shown to induce actin polymerization (Qi et al., 2009). Three of the consistently up-regulated genes (MSMO1, HMGCS1 and IDI1) in suspension HEK293 compared to adherent encode enzymes that have direct roles in the cholesterol biosynthesis pathway (Mazein et al., 2013). Methylsterol monooxygenase 1 (MSMO1), an endoplasmatic reticulum transmembrane protein, and hydroxymethylglutaryl-CoA synthase (HMGCS1), located in the cytoplasm, are catalyzing steps in the cholesterol synthesis pathway, where-as isopentenyl-diphosphate delta-isomerase (IDI1) catalyze the isomerization of one the cholesterol building blocks, isopentenyl diphosphate. Moreover, two additional genes (NPC1L1 and INSIG1) found up-regulated in suspension cells compared to adherent, are also associated with cholesterol metabolism by various processes. The NPC1L1 gene encodes a transporter protein (NPC1-like intracellular cholesterol transporter 1) required for cholesterol absorption in the intestine and is also suggested to play an important role in intracellular cholesterol trafficking between vesicles (Howles and Hui, 2012). Insulin-induced gene 1 protein (INSIG1) is a negative regulator of cholesterol synthesis and important for cholesterol homeostasis (Dong et al., 2012) and knockout of INSIG1 has previously been shown to result in cholesterol accumulation (Engelking et al., 2005). As this gene was found to be upregulated in suspension cells compared to adherent HEK293, this would have the reverse impact on the cholesterol biosynthesis pathway in suspension cells compared to the above-mentioned MSMO1, HMGCS1 and IDI1 genes, leading to a reduction of the cholesterol content of the cell. Notably, a lower cholesterol biosynthesis in suspension cell lines compared to the original HEK293 strain was indeed predicted using IPA (**Supplementary Figure 7**). It should however be noted that this prediction does not take into consideration the effect of INSIG1, instead the predicted reduction in cholesterol biosynthesis in suspension cells compared to the HEK293 cell line is a result of downregulation of SC5D (lathosterol oxidase).

From the transcriptomic data of these different HEK293 cell lines, differences between adherent and suspension cells were found to be related to differences in the metabolism of lipids in general and cholesterol specifically. This emphasizes a potentially important role of cholesterol and lipid biosynthesis in the transition from adherent to serum free suspension growth of HEK293 cell types. From a bioprocess perspective, differences in intracellular cholesterol synthesis and metabolism are of interest with regards to the secretory capacity of a cell line since cholesterol content of a cell can impact its productivity. Previous findings has shown that cholesterol is essential for ER to Golgi transport within the secretory pathway (Ridsdale et al., 2006) and that secreted productivity of CHO cells increases upon elevated intracellular cholesterol levels, through silencing of INSIG1, possibly due to increasing the volume of the Golgi compartment (Loh et al., 2017). It would therefore be of interest to gain further knowledge about the cholesterol content and distribution within HEK293 cell lines and potentially evaluate if this this pathway can be a target for enhanced bioproductivity without having a deleterious impact on suspension growth or cell morphology.

### Transcriptomic and genomic results indicated that suspension adaptation of HEK293 does not follow the epithelial to mesenchymal transition

Moreover, 4 of the 38 genes (ID1, SMAD7, TXNIP and LOX) that were consistently differentially expressed between adherent and suspension HEK293 have previously been annotated to play a role in epithelial to mesenchymal transition (EMT) (Zhao et al., 2015), the event where stationary epithelial cells lose their cell-cell adhesion and change into motile and invasive mesenchymal cells (Yang and Weinberg, 2008). However, when evaluating the expression of common markers for mesenchymal and endothelial phenotypes as well as predicting the outcome of the EMT pathway using ingenuity pathway analysis (IPA), the parental HEK293 strain showed the most mesenchymal-like phenotype whereas suspension cell lines were predicted to have reduced transition from epithelial to mesenchymal phenotype compared HEK293 (**Figure S4**). These results, along with above-mentioned details regarding copy number variation of the TLE4 and DACH1 genes, indicate that the suspension adaptation of HEK293 lineages does not follow the EMT pathway. However, suspension cells were predicted to have a higher degree of disruption of adherence junctions compared to the parental HEK293 strain (data not shown), which may be important for the suspension cell phenotype.

### Identification of five genes with potential key roles in differences between adherent and suspension human cell lines

Altogether nine of the previously identified 38 genes (LOX, ID1, ADAMTS1, ZIC1, KCNMA1, DHRS3, RARG, COL4A6 and ARRDC4) were shown to have significantly different expression levels also in an extended validation of the genes in a set of 63 human cell lines from the HPA database (Uhlen et al., 2017)(**Figure 6**). However, in the case of four of the genes (ADAMTS1, KCNMA1, COL4A6 and ARRDC4) the direction of expression between adherent and suspension cells was reversed compared to the pairwise comparison of HEK293 cell lines only (**Figure 5B**). The remaining five genes (LOX, ID1, ZIC1, DHRS3 and RARG) showed a consistent expression profile between adherent and suspension cells compared to the results presented in **Figure 5B**, suggesting a key role of these genes in the morphologies of adherent and suspension cells. ID1 belongs to the TGF-beta signaling pathway and up-regulation of this gene, as found in adherent cells compared to suspension cell lines, has been associated with the mesenchymal-to-epithelial transition (ID1) (Stankic et al., 2013). Consistently, ID1 silencing has also been shown to significantly reduce adhesion of neural stem cells (Tan et al., 2012) and conversely, increased ID1 expression in epithelial cells has been related to increased adhesion (Qiu et al., 2011). In addition, lysyl oxidase (LOX), an enzyme responsible for the covalent cross-linking between elastin and collagen in the extracellular matrix, has been shown to be important for cell-matrix adhesion formation, supporting the adherent phenotype of adherent cells but is also associated with cell invasion and induction of EMT (Payne et al., 2005)(Schietke et al., 2010). Besides the EMT-related genes, the additional three genes (RARG, DHRS3 and ZIC1), consistently up-regulated in adherent cells compared to suspension cell lines, have previously been associated with cell adhesion through the retinoic acid signaling pathway. Indeed, retinol metabolism was found to be down-regulated in suspension cells also in the metabolic gene set analysis. The RARG gene, which is upregulated in adherent cells compared to suspension cells, encodes the retinoic acid receptor gamma that controls the expression of genes involved in cell adhesion and plays an important role in promoting cell adhesion in mouse embryonic fibroblasts (Al Tanoury et al., 2013)(Kelley et al., 2017). Furthermore, the transcription factor ZIC1 has been associated with the upregulation of genes associated with retinoic acid signaling and adhesion (Cornish et al., 2009; Gan et al., 2011) whereas retinoic acid-inducible dehydrogenase reductase 3 (DHRS3) is an enzyme important for retinoic acid homeostasis and acts a negative regulator of retinonic acid synthesis. This enzyme has a weak catalytic capacity to convert retinaldehyde to retinol, which results in reduced retinoic acid levels, unless co-expressed with RDH10, which induces the full capacity of the DHRS3 enzyme, resulting in decreased retinoic acid synthesis (Adams et al., 2014). Furthermore, signaling of retinonic acid receptors only in the absence retinoic acid is associated with increased adhesion in mouse embryonic fibroblasts (Al Tanoury et al., 2013). The level of RDH10, along with DHRS3, was significantly lower in the HEK293 suspension cells compared to adherent (**Table S5**), suggesting more efficient catalytic activity of the DHRS3 enzyme in the adherent cell lines, which may lead to retinoic acid signaling in the absence of retinoic acid, resulting in a more adherent phenotype in these cell lines.

### Conclusions

Our study has outlined the genomic, transcriptomic and metabolomic variations between six industrially relevant HEK293 cell lines, in an attempt to improve the understanding of their respective differences in phenotype. We report a selective pressure to develop certain expression profiles during the evolution and continuous cultivation, evidenced by the numerous genes and pathways detailed here. The key common changes between HEK293 and its progeny cell lines involve in particular cell membrane proteins and processes related to cell adhesion, motility and the organization of various cellular components such as the cytoskeleton and extracellular matrix. In addition, changes associated with differences between adherent and suspension cell growth in particularly involve changes in cell adhesion protein expression, cholesterol metabolism and a set of six key genes (RARG, ID1, ZIC1, LOX and DHRS3) with potentially key roles in the differentiation between the two groups. These results could be of importance when pursuing further cell line engineering or bioprocess optimization of these and other human cell lines.

## Acknowledgements

This work was supported by the Knut and Alice Wallenberg Foundation, AstraZeneca, SSF, Vinnova and the Novo Nordisk Foundation (grant no. NNF10CC1016517).

## Author Contributions

Conceptualization, R.F., P.V., M.U., J.N., and J.R; Methodology, M.M., R.S. and J.R. Formal analysis, M.M., R.S., and M.L.; Investigation, M.M., R.S., and M.L., Writing – Original Draft, M.M, R.S., M.L., and J.R.; Writing – Review & Editing, M.M., M.L., D.H., J.N., and J.R.; Visualization, M.M., R.S., and M.L., Supervision, R.F., P.V., D.H., T.S., M.U., J.N., and J.R., Funding Acquisition, R.F., P.V., M.U. J.N, and J.R.

## Declaration of interests

The authors declare no competing interests

---

Table S1. Transcriptomic data (TPM values) of all genes in six HEK293 cell lines

Table S2. Genes with full or partial copy number gain or loss (>1/<−1 log2 foldchange) in HEK293 progeny cell lines compared to the parental HEK293. Related to Figure 2.

Table S3. High and moderate impact SNPs common and unique to all progeny cell lines compared to the parental HEK293.

Table S4. Results of differential expression analysis between progeny cell lines compared to HEK293 and intersects between results. Related to Figure 3 and Figure 5.

Table S5. Results of differential expression analysis between suspension and adherent progeny HEK293 cell lines. Related to Figure 4.

Table S6. List of differentially expressed genes (padj <0.01) in suspension compared to adherent HEK293 cell line. Related to Figure 4.

Table S7. Full results of metabolic gene set analysis. Related to Figure 4.

Table S8. Normalized counts of 38 differentially expressed genes between suspension and adherent HEK293 cell lines in HPA cell lines. Related to Figure 6

